# The sensitivity of transcriptomics BMD modeling to the methods used for microarray data normalization

**DOI:** 10.1101/781567

**Authors:** Roman Mezencev, Scott Auerbach

**Affiliations:** National Center for Environmental Assessment, US EPA, Washington DC, USA; National Institute of Environmental Health Sciences, NIH, Research Triangle Park NC, USA

**Keywords:** Toxicogenomics, transcriptomics, microarray, normalization, gene expression, dose-response modeling, BMD, benchmark dose, POD, point of departure, RMA, GCRMA, MAS5.0, PLIER, PLIER16

## Abstract

Whole-genome expression data generated by microarray studies have shown promise for quantitative human health risk assessment. While numerous approaches have been developed to determine benchmark doses (BMDs) from probeset-level dose responses, sensitivity of the results to methods used for normalization of the data has not yet been systematically investigated. Normalization of microarray data converts raw hybridization signals to expression estimates that are expected to be proportional to the amounts of transcripts in the profiled specimens. Different approaches to normalization have been shown to greatly influence the results of some downstream analyses, including biological interpretation. In this study we evaluate the influence of microarray normalization methods on the transcriptomic BMDs. We demonstrate using *in vivo* data that the use of alternative pipelines for normalization of Affymetrix microarray data can have a considerable impact on the number of detected differentially expressed genes and pathways (processes) determined to be treatment responsive, which may lead to alternative interpretations of the data. In addition, we found that normalization can have a considerable effect (as much as ∼30-fold in this study) on estimation of the minimum biological potency (transcriptomic point of departure). We argue for consideration of alternative normalization methods and their data-informed selection to most effectively interpret microarray data for use in human health risk assessment.

## Introduction

Whole-genome expression data generated by microarray studies are a promising resource for human health risk assessment. Analysis of these data can provide insight into mechanisms of biological processes, enable prediction of adverse outcomes of chemical exposures, and support estimation of points of departure (PODs) for derivation of toxicity values (reviewed in [1]).

Genomic dose-response studies (GDRS) done in an *in vivo* setting have been shown to identify gene set (e.g., pathway)-level PODs that approximate those identified using much more resource intensive guideline toxicity assessments [2,3]. In addition, mechanistic interpretation of GDRS data can yield a deeper understanding of molecular effects produced by tested substances in a manner that can support Adverse Outcome Pathway (AOP) development and human relevance determination [4].

One approach to analysis of GDRS data is embodied in the BMDExpress software [5] which carries out a three-step process to identify gene set-level biological potency estimates referred to as benchmark dose (BMD) values. The three-step process includes (i) identification of probe sets (genes) responsive to chemical exposure, (ii) dose response model fitting of treatment responsive probe sets, and (iii) summarization of gene-level BMDs as gene set (e.g., pathway) level BMDs. Pathway level BMD and their lower confidence limits (BMDLs) are subsequently interpreted in the context of the lowest doses at which biological changes occur (i.e., biological effect point of departure [BEPOD] and biological point of departure lower bound [BEPOD/L], in the case of the lowest BMD and associated BMDL, respectively).

At each level of the analysis, parameter selection (e.g., choosing a minimum fold change) is made that can dramatically impact the amount of information that is carried through the analysis. For the assessment discussed here we have used a set of parameters identified by the National Toxicology Program (NTP) [6].

Before BMD modeling of microarray expression data can take place, the raw fluorescence signals must be processed. This processing, frequently referred to as “normalization”, mathematically transforms raw signals to expression estimates that are proportional to the amounts of corresponding transcripts in the profiled specimens. The need for normalization relates to a complex set of processes that can introduce non-biological variability, along with complex relationships between input quantities of mRNAs and signal intensities. These processes include, but are not limited to reverse transcription, labeling and hybridization on microarrays. Normalization of raw data from Affymetrix microarrays has been the focus of a remarkable amount of research due to the significant effect it can have when interpreting microarray data and the popularity of the Affymetrix platform. Numerous normalization methods are currently available, including MAS5.0.0, RMA, GCRMA, and PLIER. Each of these methods includes background adjustment (separates the specific signal from the non-specific signal), normalization and probe summarization steps [7]. Due to differences in assumptions underlying each of these methods[8], selection of normalization methods can significantly impact results of downstream analyses, such as identification of differentially expressed genes [9], clustering of genes or specimens [10], development of gene expression-based classifiers and building gene networks [11]. Previously published GDRS employed the most commonly used method for normalization of Affymetrix microarrays RMA (Robust Multi-array Average) [3,12-15]. Here we investigate the influence of different normalization methods on genomic dose-response modeling.

In this article we demonstrate the influence of different normalizations of expression microarrays on the findings from 5-day *in vivo* genomic dose-response studies of crude 4-methylcyclohexanemethanol (crude MCHM) [16], neat 4-methylcyclohexanemethanol (MCHM) [16], N,N-dimethyl-p-toluidine (DMPT) [17] and p-toluidine [17] in liver tissues, and propylene glycol phenyl ether (PPH) [16] in kidney tissues of orally-exposed rats. Our results indicate that normalization remarkably impacts the number of detected differentially expressed genes and responsive gene sets. In addition, we demonstrate that the different normalization methods can lead to changes in model fitting, which alter individual probe set BMD values. Estimates of minimum biological effect potency (BEPOD/L) were found to be robust to the effects of different normalization methods for some but not all chemicals. In some cases, potentially impactful differences between BEPOD/L values were observed with different normalization methods, hence this variable should be considered carefully when performing genomic dose response analysis.

## Materials and Methods

### Gene Expression Data

All expression data used in this study were generated by profiling liver specimens of rats using the Affymetrix Rat Genome 230 2.0 Array that contains more than 31,000 probe sets analyzing transcripts from 15,575 annotated unique genes (NetAffx Annotation File, Release 36; URL: https://www.thermofisher.com/order/catalog/product/900505).

Raw expression data for livers of male Harlan Sprague Dawley or male F344/N rats exposed orally for 5 days to crude 4-methylcyclohexanemethanol (crude-MCHM), or neat 4-methylcyclohexanemethanol (MCHM), and for kidneys of rats exposed orally to propylene glycol phenyl ether (PPH) were accessed from the Gene Expression Omnibus (GEO) as series GSE75655, GSE75657 and GSE75656, respectively. Hepatic transcriptomics data for F344/N rats exposed orally to N,N-dimethyl-p-toluidine (DMPT) or p-toluidine for 5 days [17] were accessed from the GEO as series GSE100502. All .cel files available for each replicated exposure level and vehicle controls were included in this analysis.

Crude MCHM represents a mixture of six components in addition to the major component 4-methylcyclohexanemethanol (MCHM) used as a coal cleaning liquid. Propylene glycol phenyl ether (PPH) is an industrial chemical used as a latex coalescent and a solvent for textile dyes. Toxicological significance of MCHM and PPH is associated with a recent spill and large scale contamination of drinking water in West Virginia [18]. N,N-dimethyl-p-toluidine (DMPT) is an accelerant for methyl methacrylate monomers in medical devices that has been shown to induce liver carcinogenesis in male and female F344/N rats and B6C3F1 mice in a 2-year oral exposure study. p-toluidine, structurally-related to DMPT, is reportedly a liver carcinogen in mice [17].

### Data processing

The raw fluorescence signals was processed using the following methods: MAS5.0.0 (Microarray Affymetrix Suite version 5.0), RMA (Robust Multichip Analysis), GCRMA (GeneChip Robust Multichip Analysis) and PLIER (Probe Logarithmic Intensity Error Estimation) [7,19]. Raw data in .cel file format were imported into Expression Console Build 1.3.1.187 (Affymetrix, Santa Clara, CA, USA) and processed using MAS5.0 and RMA methods with default configurations. Processing using PLIER method was performed with PM-MM background correction and quantile normalization. Processing using GCRMA method was implemented in R version 3.5.1 (https://www.R-project.org) using the “GCRMA: Background Adjustment Using Sequence Information” R package version 2.52.0 (Bioconductor version 3.7). PLIER16 values were calculated from PLIER values by adding 16 to each signal intensity value as a simple variance-stabilizing transformation. Low-quality microarrays were identified by visual inspection of the Relative Log Expression (RLE) boxplots generated by Expression Console (Affymetrix). RLE values are calculated for each probeset as the ratio between expression of this probeset in a given microarray and the median expression of this probeset across all the arrays in the dataset [20]. Low-quality microarrays were removed from the datasets and the raw data were re-normalized by GCRMA, RMA and PLIER methods (MAS5.0 uses a per-chip approach and re-normalization was not needed). Normalized data used in this study for BMD modelling are available at the URL: https://catalog.data.gov/organization/epa-gov

### Differential expression analysis and transcriptomics BMD modeling

Normalized data were imported into BMDExpress for Windows, version 2.20 build 0167 BETA (https://github.com/auerbachs/BMDExpress-2/releases) [21]. For all chemicals, differentially expressed genes were determined from normalized datasets with all probesets included. To examine the influence of non-informative probesets, the analysis was also performed following removal of the probesets that showed “Absent” MAS5.0 absolute detection calls across all specimens from MAS5.0 and PLIER16-normalized data (MAS5.0_noA calls and PLIER16_noAcalls methods).

Differentially expressed probesets were detected using the Williams trend test (p<0.05; absolute fold change ≥1.5). Benchmark response of 1 SD was used for each feature. Best models among linear, 2nd degree polynomial, Hill, power and exponential models (degrees 2-5) were selected based on the lowest AIC. For more details on parameters used in BMDExpress, see Supplemental File. Gene set-level BMD values were determined by mapping probes that met BMD filtering criteria (Supplemental File) to GO: Biological Process ontologies and Reactome Pathways [22]. The most sensitive GO:BP (Gene Ontology Biological Process) or Reactome pathways were identified as those with lowest median BMD or BMDL values calculated from all mapped probesets and reported as BEPOD or BEPOD/L, respectively.

### Visualization and statistics

Comparison of differentially expressed genes was visualized using Venn diagrams (http://bioinformatics.psb.ugent.be/cgi-bin/liste/Venn/calculate_venn.htpl). All other plots were produced using TIBCO Spotfire Analyst version 7.8.0 (TIBCO Software Inc, Palo Alto, CA 94304, USA). Agreement between probe-level BMD or BMDL values determined by different normalization methods was assessed by Pearson correlation using Partek Genomics Suite version 7.18 (Partek Incorporated, St. Louis, MO 63141, USA). Statistical significance of differences among probeset-level best model fit BMD and BMDL values for PPH determined by 7 normalization methods were tested using Kruskal-Wallis test and the differences were considered statistically significant for two-tail p-values<0.05. Most sensitive GO:BP gene sets were summarized for visualization using Revigo tool [23].

## Results

### Assessment of Normalization

We explored the influence of 7 different microarray normalization methods and two different collections of gene sets on estimation of BEPOD/L values for five test articles using public data generated by genomic dose-response studies. Methodological differences between 7 used normalization methods are summarized in (Supplemental file, Table 1).

Visual inspection of RLE plots for raw MCHM and PPH data identified low-quality microarrays that differed from other microarrays in medians and distributions of their RLE values. These microarrays were removed from datasets and differentially expressed genes (probe sets) and BMD values were determined from the remaining expression data (Supplemental File, Figures 1-10)

### Effect of normalization on the identification of differentially expressed probe sets and active gene sets

Different normalization methods identified remarkably different numbers of differentially expressed probe sets (DEPSs). The highest numbers of DEPSs for most chemicals were identified by using MAS5.0 and PLIER methods, while PLIER16 and PLIER16-noAcalls identified considerably less probe sets corresponding to differentially expressed genes (Figure 1A). Likewise, GCRMA method produced more DEPSs than RMA with the single exception of p-toluidine (Figure 1A).

**Figure 1.**
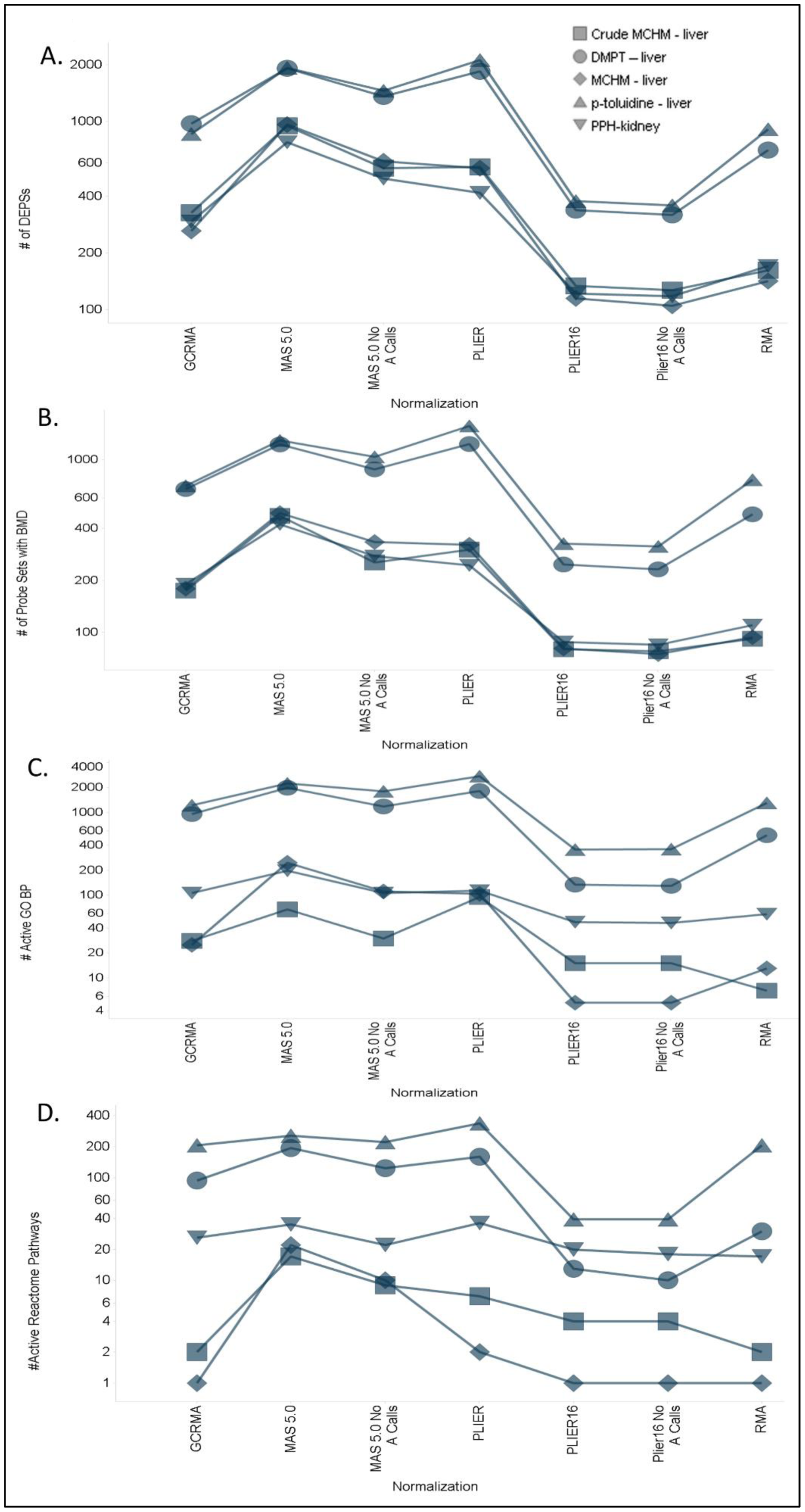
Effect of different normalizations on the number of (A) differentially expressed probe sets (DEPSs), (B) probe sets with acceptable BMD estimates, (C) active GO Biological Processes and (D) active Reactome Pathways.

The use of MAS5.0 and PLIER also produced the highest numbers of DEPSs with acceptable model fits (AMFs; i.e. DEPSs where the global goodness of fit p-value >0.1 and the BMDU/BMDL ratio <40), which would be therefore included in the gene set BMD analysis (Figure 1B). Consistent with this observation, these two methods produced highest numbers of GO:BP and Reactome Pathways, from which gene-set-level BMDs and ultimately BEPOD values could be determined (Figure 1C and 1D)). In contrast, RMA, GCRMA, PLIER16 and PLIER16_noAcalls methods generally identified fewer DEPSs for which acceptable model fits, and therefore all identified fewer “active” GO:BP and Reactome pathways that could be used for estimation of BEPOD (Figure 1C and 1D).

Comparison of overlaps among DEPSs with acceptable model fits across the 7 normalization methods was also performed for all data sets. In all chemical-organ sets, the percent of overlapping DEPSs with AMFs was quite small (<10%) when all normalizations were compared, suggesting that normalization is likely to have a large impact on the qualitative interpretation (e.g., mode of action, AOP assessment) of the toxicogenomic effects. To obtain a sense of how great the effect normalization has on the identification of DEPS with AMF, an “intersection” set for all normalizations per treatment-chemical pair was identified and used as a comparator for each normalization in a given chemical-treatment set (Figure 2A). This analysis showed that normalization methods such as MAS5.0 and PLIER consistently identified a greater number of DEPSs with acceptable model fits compared with other normalization methods. To more accurately quantify the relative increase and DEPSs with AMFs, a fold increase over intersection for each normalization in a chemical-organ pair was calculated (Figure 2B). MAS5.0 (median ∼9.5) and PLIER (median ∼7) showed the greatest fold increase in DEPSs with BMDs whereas PLIER16 normalizations showed the smallest fold increases of intersection (Figure 2B). These findings are consistent with the relatively larger number of active GO:BP and Reactome pathways that are identified when using MAS5.0 and PLIER normalized data (Figure 1C and D). In addition, these results are consistent with analysis of overlaps among differentially expressed genes identified by different normalization methods (Supplemental file, Figure 11).

**Figure 2.**
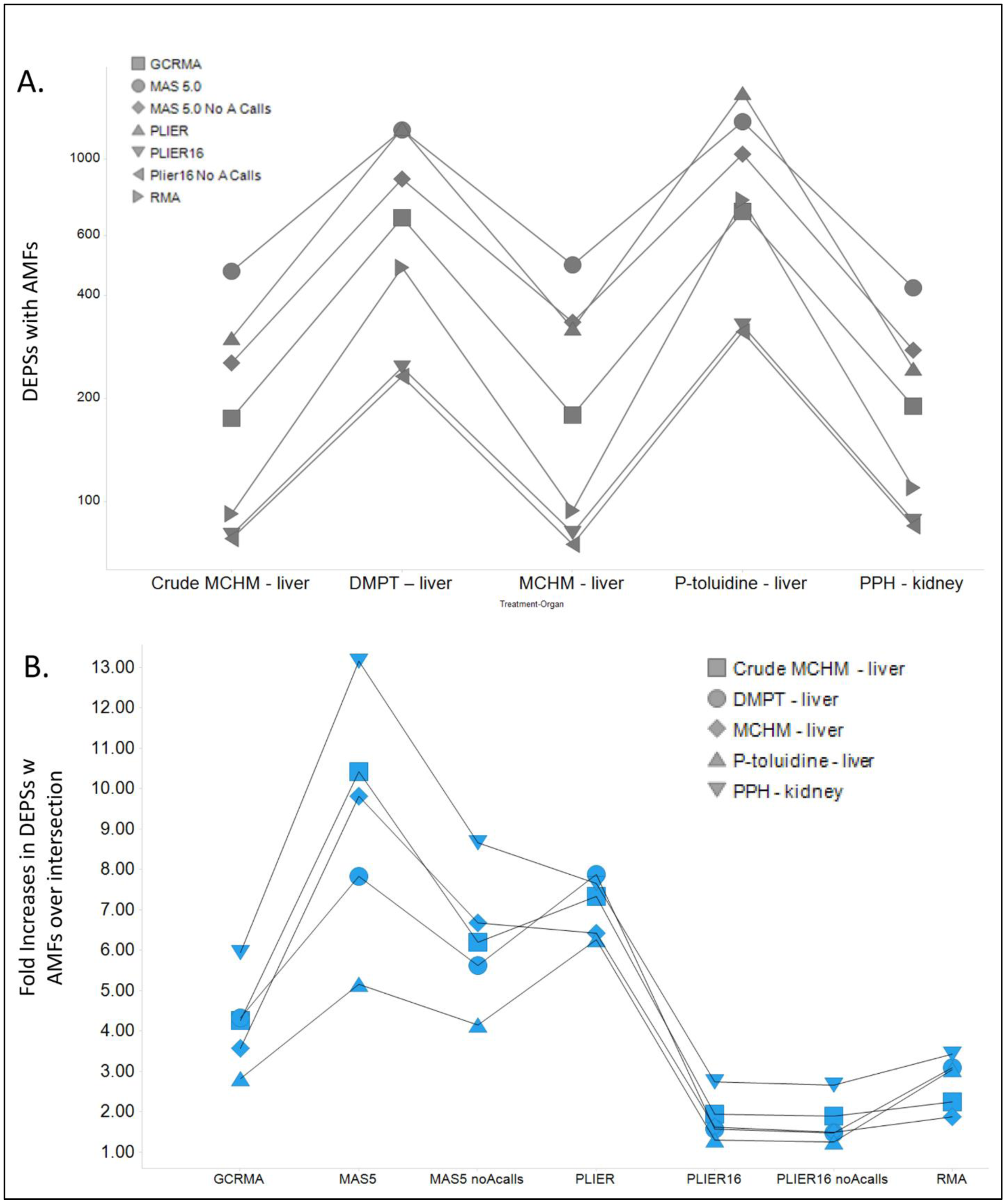
(A) Effect of different normalizations on the number of differentially expressed probe sets (DEPSs) with acceptable model fits (AMFs). (B) Fold increases in DEPSs over an intersection of all normalizations for any individual chemical-treatment pairs.

### Effect of normalization on the distribution of probeset BMD and BMDL

Distributions of probeset-level BMD and BMDL values showed differences across normalization methods (Figure 3A and B). For example, differences among distributions of best model fit BMD and BMDL values for PPH were found statistically significant (Kruskal-Wallis p<0.0001 for BMD and BMDL). Median probeset-level BMD value for MAS5.0 method (BMD=637.4 mg/kg-day) was higher than median BMD values determined by all other normalization methods (range: 341.6-583.9 mg/kg-day); however, the lowest probeset-level BMDs for PLIER16 and PLIER16_noA (71.1 mg/kg-day) were found substantially higher than corresponding values for MAS5.0 and other normalization methods (range: 0.36-19.3 mg/kg-day) (Figure 3A). Notably, median probe-set level BMD values were highest for MAS5.0 normalization across all chemical-treatment pairs (Figure 3C). Similar manifestations are present in the gene set analysis where the GO:BP median BMD value distributions were higher when using MAS5.0 and generally lowest with PLIER16 (Figures 3D- and 3E)

**Figure 3.**
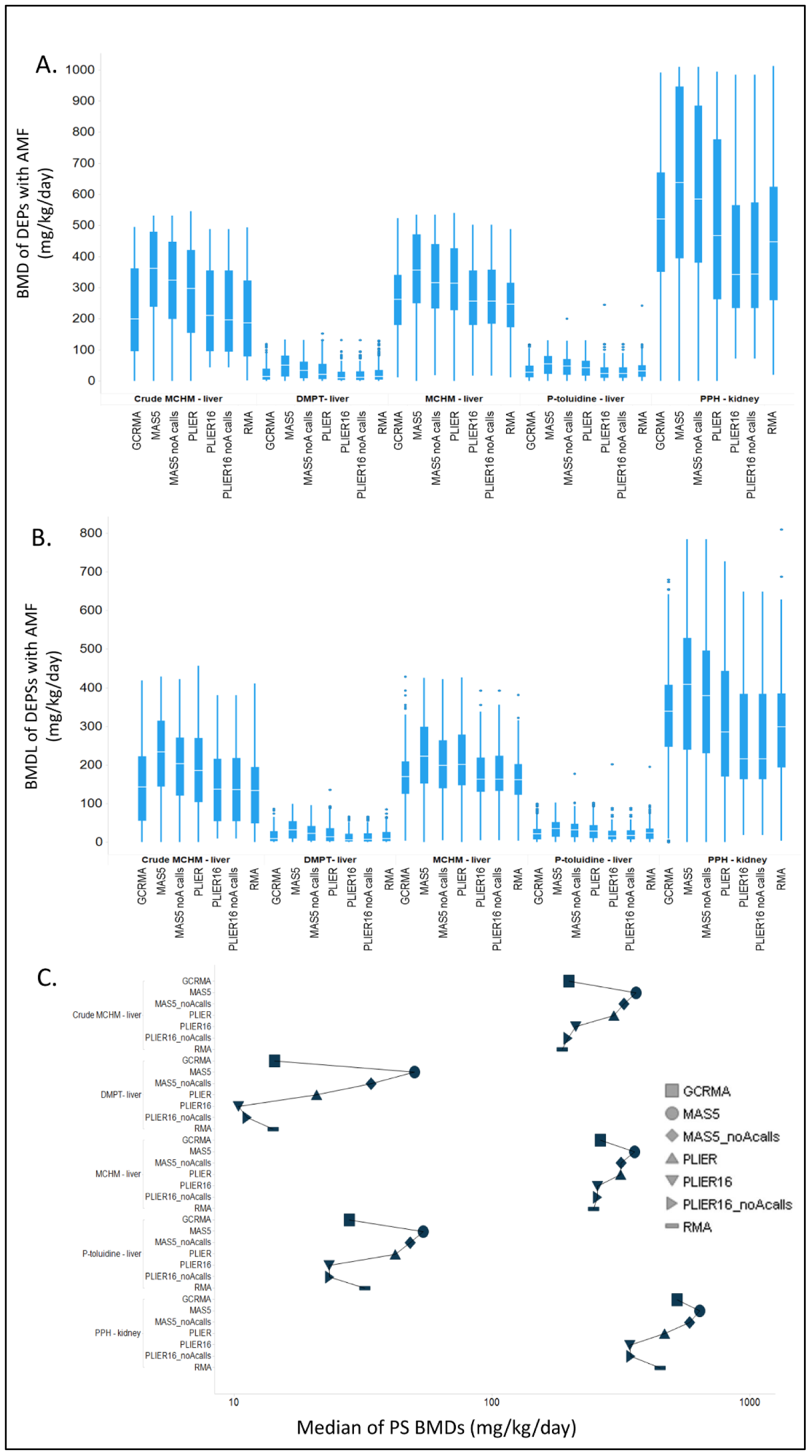
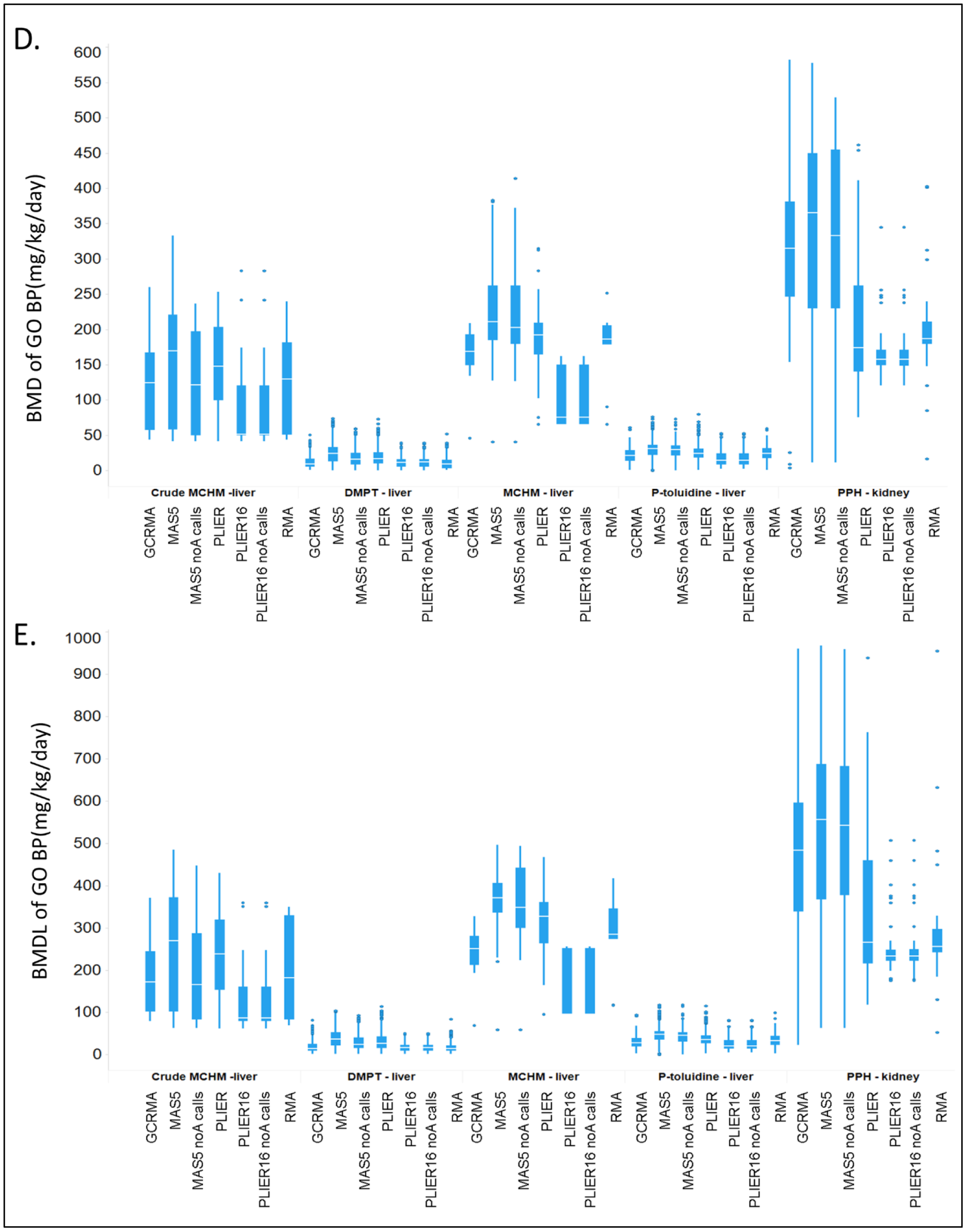
Impact of normalization on the overall distribution of probe set-level BMD (A) and BMDL (B) values and GO:BP median BMD (D) and BMDL (E) values in each of the 5 experiments.

### Effect of normalization on the lowest gene set BMD and BMDL (BEPOD and BEPOD/L) determination

Selection of different normalization methods influenced determination of lowest median gene set-level BMD (BEPOD) and BMDL (BEPOD/L) for different chemicals to a different extent. Within a gene set type for any one treatment, the BEPOD/L variation ranged from 1.1 fold (Crude MCHM, Reactome BMDL) to up to 30.4 fold (P-toluidine, GO:BP BMDL) due to different normalizations (Figure 4 and Supplemental table 2). The mean and median variations in BEPOD/L values for a chemical treatment-gene set pair due to different normalizations were 6.2- and 3-fold, respectively. In general, MAS5.0 normalization tended to yield the lowest BEPOD/L values and PLIER16 the highest (Supplemental figure 12). Further, GO:BP gene sets tended to produce lower BEPOD/Ls compared to the Reactome Pathway set, likely due to the relatively larger number curated gene sets included in the GO:BPs.

**Figure 4.**
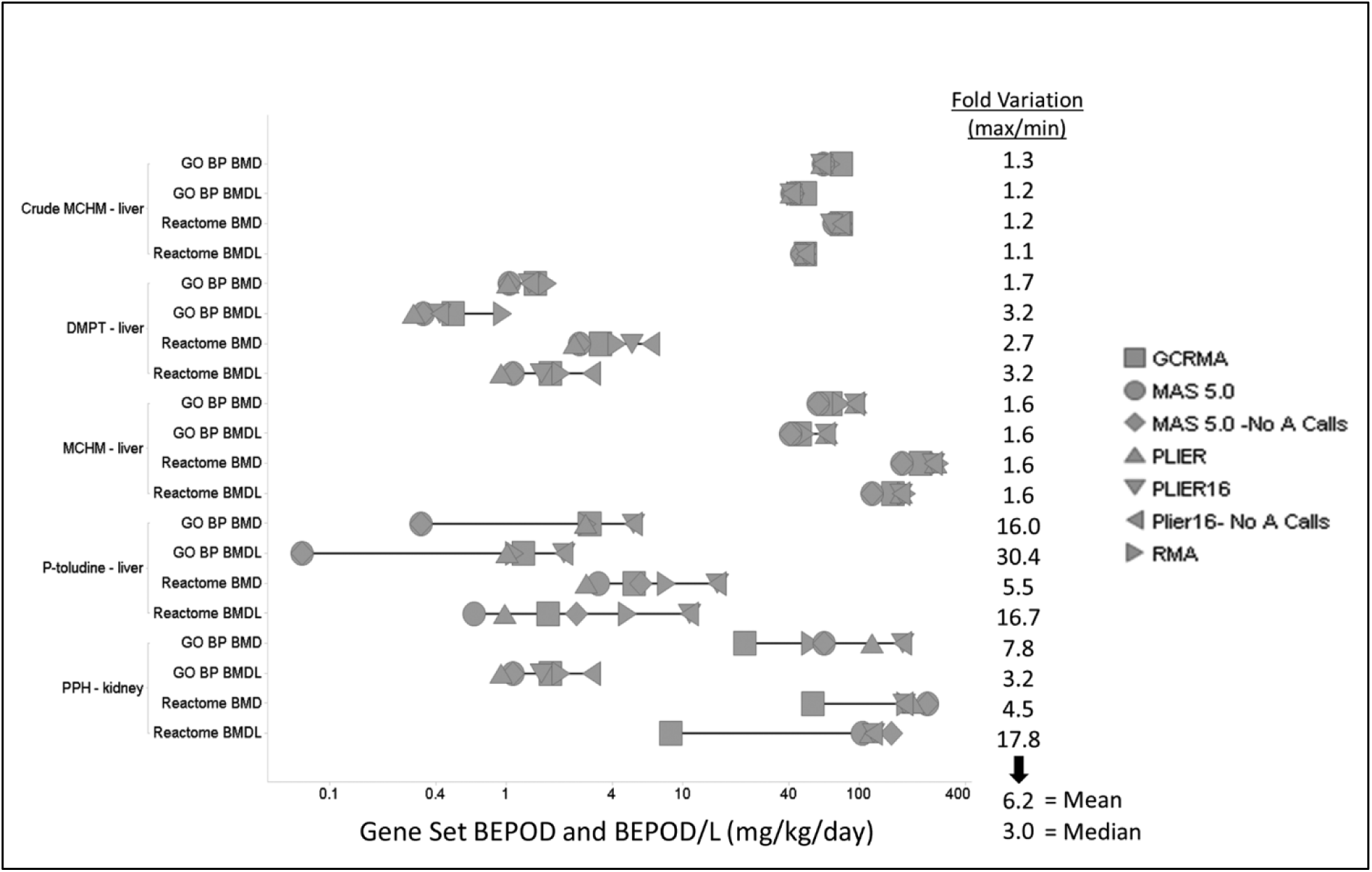
Effect of normalization on the lowest median BMD (BEPOD) and BMDL (BEPOD/L) values for GO:BP and Reactome pathways. The distribution of BEPOD and BEPOD/L values across 7 normalizations are shown for each of the 5 experiments. Fold variation: max BEPOD to min BEPOD (or max BEPOD/L to min BEPOD/L) ratio. Mean and median fold variations are shown for BEPOD/L vales.

### Effect of normalization on the identity of the gene sets with the lowest BMD and BMDL (BEPOD and /L)

Influence of normalization methods on the identity of the most sensitive GO Biological Processes and Reactome pathways varied across datasets (Figure 5A and B). For instance, the same three most sensitive GO:BP gene sets (GO:0002933, GO:0070988, and GO:0070989) were found for all normalization methods in the case of the MCHM data set (Figure 5A and Supplemental table 3). In contrast, a variety of different GO:BP gene sets were identified as the most sensitive when using different normalizations of the p-toluidine dataset. While “MAS5.0_noA calls” and “PLIER16_noA calls” detected the same most sensitive pathways as their corresponding parental methods MAS5.0 and PLIER16, other normalization methods identified seemingly biologically unrelated most sensitive GO:BP gene sets such as, e.g. GO:00043588 “Skin development” (RMA), GO:0044246 “Regulation of multicellular organismal metabolic process” (GCRMA), GO:0007040 “Lysosome organization” (MAS5.0), and GO:0048266 “Behavioral response to pain” (PLIER) (Figure 5C). Further, median BMD values corresponding to these pathways were also appreciably different (Supplemental table 2).

**Figure 5.**
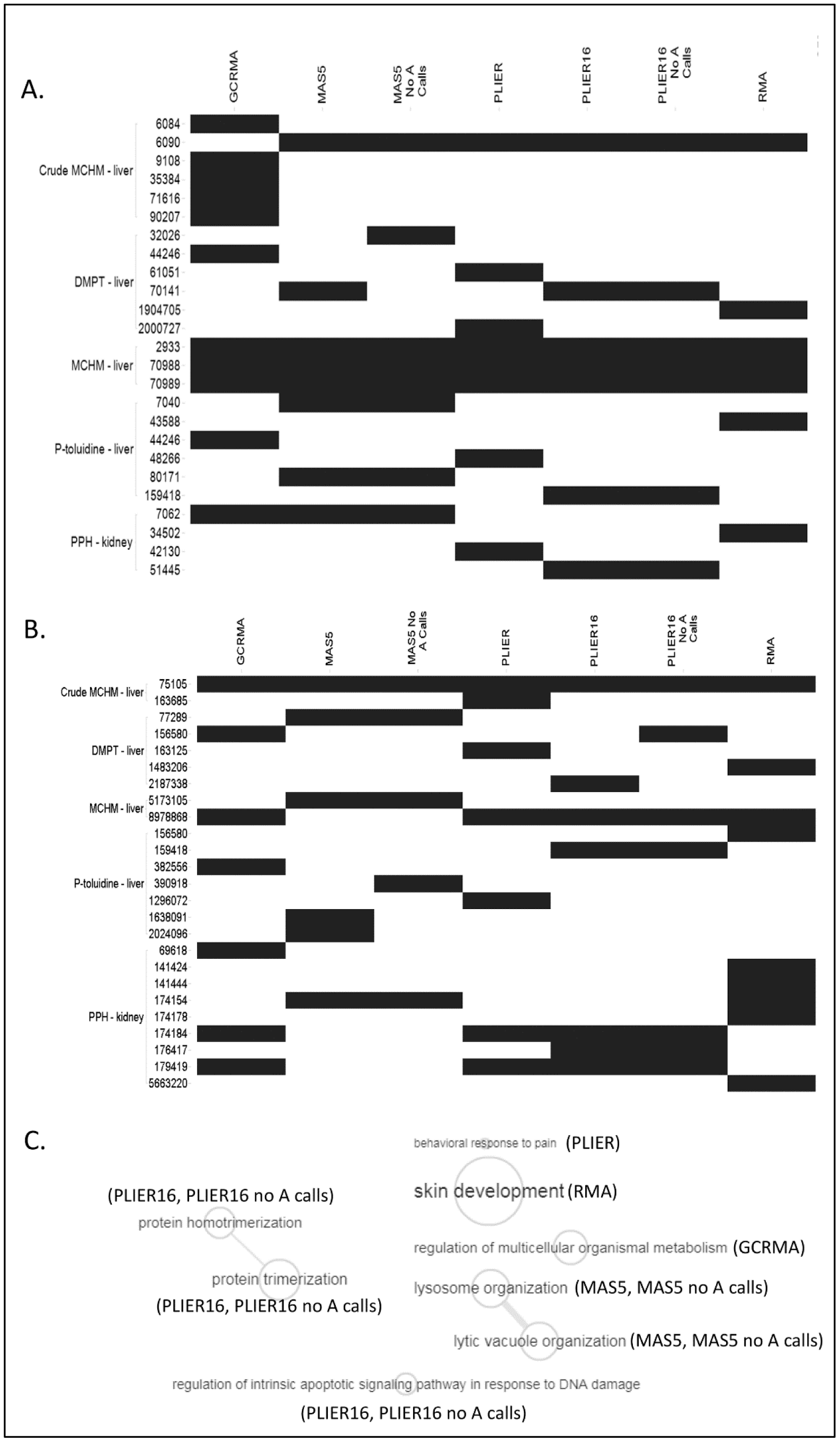
GO Biological Processes (A) and Reactome Pathways (B) that were identified as most sensitive when different normalizations were employed. A black cell indicates that the gene set was identified as the gene set with the lowest median BMD value. (C) A semantic similarity visualization of the GO:BP terms with lowest median BMD values generated using REVIGO (allowed similarity =0.9; database with GO term sizes: Rattus norvegicus; similarity measure: SimRel).

### Agreement of Gene Set BMD and BMDL values across normalizations

To determine the effect of normalization on overall agreement of the BMD values, Pearson correlation between both probe set BMD values and median BMD values for GO:BP and Reactome pathways in each of the five experiments was performed. In nearly all cases, BMD values from probe sets (PS) or gene sets exhibited positive correlation, however the range of Pearson correlation coefficients (PCCs) varied from as high as 1, with average PCCs for any pair of normalizations mostly falling between 0.2 to 0.4 (Figure 6A). In most cases, PS BMDs and median BMDs had higher agreement for GO:BP than for Reactome pathways. Evaluation of individual pairings of normalizations across experiments demonstrated the MAS5.0 group of normalizations and PLIER16 group normalization show high intragroup similarity across all PSs and gene set BMDs (Figure 6B). Other pairings such as RMA-MAS5.0, PLIER16-MAS5.0, PLIER-MAS5.0 show considerably worse agreement. Notably, the agreement of BMD values from Reactome pathways was found to be more negatively affected by changes in normalization than BMD values for GO:BP gene sets.

**Figure 6.**
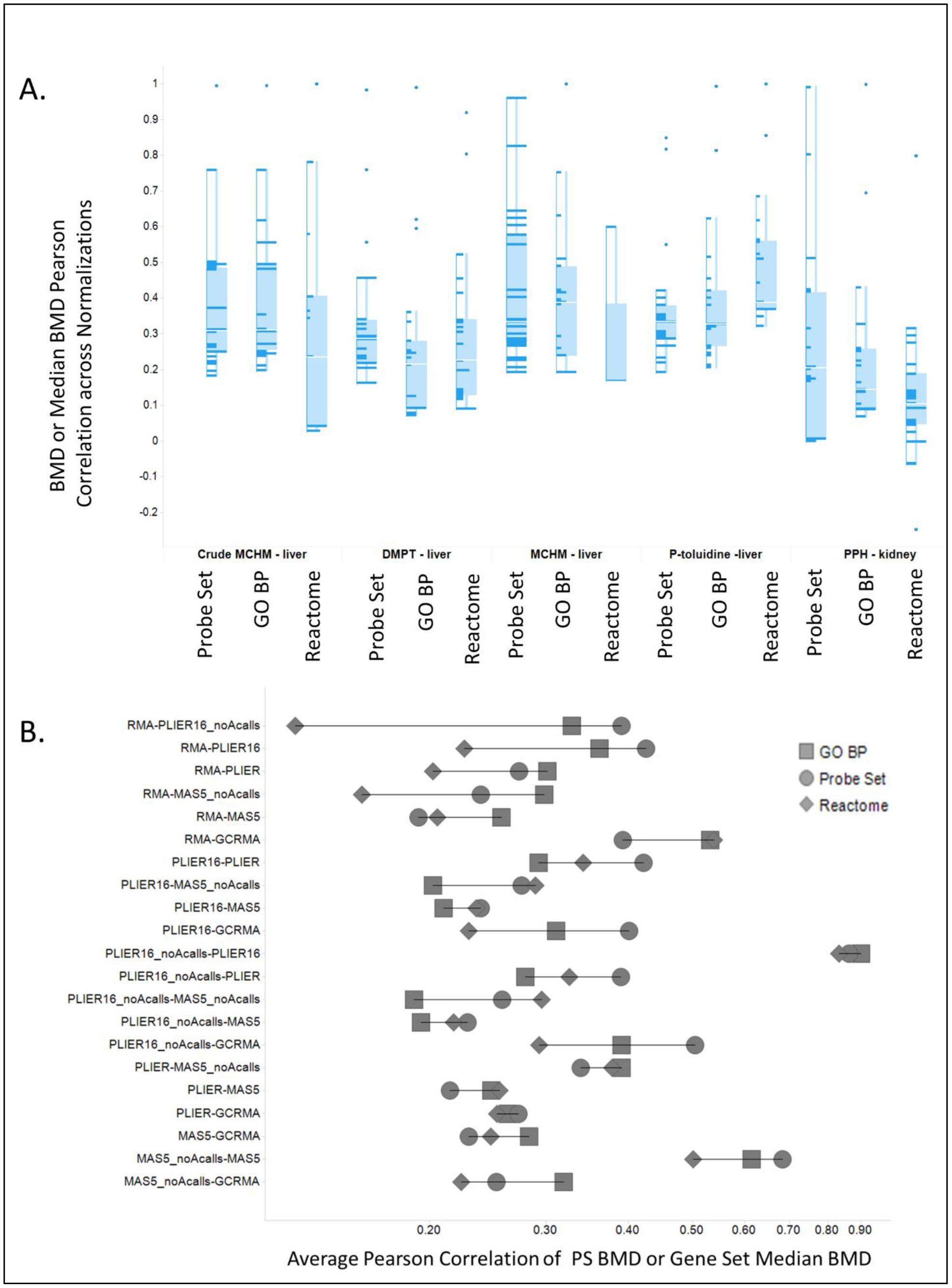
Agreement of BMD values generated by different normalizations. (A) The distribution and 95% confidence interval of the Pearson correlation coefficients (PCCs) reflecting agreement of probe set-level or gene set-level BMD values between different pairs of normalizations. For each experiment, PCCs for 21 pairs of normalizations were aggregated and presented separately for best model fit probeset level BMDs, lowest median GO:BP BMDs and lowest median Reactome pathway BMDs. (B) Average PCC values are shown for 21 different pairings.

## Discussion

Gene expression profiling has been identified as a promising method to address challenges in chemical risk assessment that have traditionally relied on data generated by time and resource consuming animal studies. This promise is especially true for short-term, in vivo, dose-response microarray studies that demonstrated potential to produce transcriptomics benchmark doses in good agreement with traditional toxicological studies [1,2].

Unlike RNA-seq where methods for analysis are still evolving, gene expression microarrays are an established tool in transcriptomics studies. Considering the relative maturity of data analysis of expression microarrays, they will likely continue to play a substantial role in whole-genome expression studies particularly in cases where the data will be used for critical decision making (e.g., in regulatory context) [1].

Analysis of microarray expression data needs to address a non-biological variability introduced by sample preparation, labeling, hybridization, fluorescence reading and other technical issues through normalization of raw microarray data. Normalization procedures attempt to remove non-biological variability from data by exploiting and enforcing known or assumed invariances of the data using different approaches [24]. Not surprisingly, numerous normalization methods have been developed for different platforms of expression microarrays [7,19]. These methods can produce considerably different results of downstream studies utilizing gene expression data, which was previously shown for some specific types of analyses, such as class discovery and gene co-expression analyses [9-11]. This study is the first to systematically evaluate the effect of microarray normalization on the transcriptomic BMD modeling. Considering the potential influence of normalization methods due to their different underlying assumptions and performance, the absence of insight on microarray normalization, especially in the context of quantitative toxicogenomics, may be perceived as a gap impairing the use of microarray data in chemical risk assessment.

The purpose of this study was to examine sensitivity of transcriptomics BMD modeling to some frequently used methods for microarray data normalization. Our results demonstrate that the use of different normalization methods produces considerably different lists of differentially expressed genes. Further, we found that the influence of microarray normalization methods on the results of transcriptomics BMD modeling is limited on some, but considerable on other datasets. We show that the fold differences between BMDL values determined from microarray data normalized by different methods can be as low as 1.1 -fold, but also as big as 30.4-fold. These differences are potentially noteworthy, because transcriptomic and apical BMD values can differ by factor of ten, when transcriptomic values are determined using RMA normalization [3]. Historically, genomic BMD analysis using Affymetrix microarrays has relied exclusively on RMA method; however, the findings presented here suggest that the appropriateness of the choice of normalization methods should likely be considered on a case-by-case basis. Consideration on a case-by-case basis instead of choosing a single method as suitable for all data reflects the fact that normalization methods differ in their fundamental assumptions, and these assumptions are satisfied by real data to varying degrees. This consideration may include behavior of normalized data (e.g., through RLE and MA plots), number of genes and gene sets that could be used for determination of BEPOD and BEPOD/L values, emphasis on sensitivity/specificity and possibly also examination of quality of probe sets that mapped to the most sensitive gene sets and subsequently projected into BEPOD and BEPOD/L values. If decision among several normalization methods cannot be made, normalizations that provide lowest BEPOD estimates for specific datasets will be interesting in the context of quantitative risk assessment as the most protective. In our study, MAS5.0 and MAS5.0_noA methods provided lowest (or next to lowest) BEPOD and BEPOD/L estimates for three datasets, GCRMA for one dataset, and all normalization methods were equivalent for one dataset.

Prior comparisons of microarray normalization methods have not produced a clear winner. While some investigators found GCRMA to perform as well as or better than other methods [25], others reported good performance of PLIER and its superiority over MAS5.0 [26], and some studies favored RMA over other methods [27]. In gene expression correlation studies, MAS5.0 reportedly outperformed GCRMA, RMA and Li-Wong methods [11]. Normalization methods differ in their precision and accuracy and the most precise methods have been shown to be generally less accurate, while more accurate methods tend to have low precision [19]. For instance, RMA offers higher precision than MAS5.0 method, which introduces high variability particularly into low-intensity probes. Nevertheless, MAS5.0 provides linear relationship between signal and transcript concentration even at low transcript concentrations when RMA method introduces bias [19], and the use of MAS5.0 alongside with detection calls substantially improves its performance to detect differentially expressed genes [28]. Furthermore, relative importance of precision and accuracy seems to depend on the purpose of transcriptomics analysis. For instance, classification and clustering problems benefit from more precise methods, because in these analyses variability accumulates from all probe sets and quickly impairs their results. While we do not attempt to propose a guideline regarding the use of microarray normalization methods, we argue that different normalization methods need to be considered as a part of quantitative toxicogenomic studies and that sensitivity of reported BEPOD and BEPOD/L values to different normalization, or justification of appropriateness of the selected normalization method need to be provided. Consequently, RMA pipeline, which has been almost exclusively used for processing of Affymetrix microarray data in quantitative toxicogenomics, should not be applied indiscriminately.

### Disclaimer

The views expressed are those of the authors and do not necessarily represent the views or policies of the U.S. Government, and they may not be used for advertising or product endorsement purposes.

## Supporting information

BMDExpress 2.2 Project File

