## Supplemental methods, tables and figures for "The sensitivity of transcriptomics BMD modeling to the methods used for microarray data normalization"

Mezencev R. and Auerbach S.

### 1. Supplemental methods

1.1 Transcriptomic BMD modeling parameters:

- Prefilter: Williams' trend test; p-value cutoff: 0.05; Filter out control genes (probes starting AFFX...); Use Fold Change Filter 1.50
- Benchmark Dose Analyses: Continuous models Hill (v. 2.18), Power (v. 2.19), Linear, Poly 2 (v. 2.21), Exp2, Exp3, Exp4, Exp5 (v. 1.11); Restrict Power ≥1; Confidence Level=0.95; Constant Variance; Maximum iterations=250; BMR=1 SD
- Best Model Selection: Lowest AIC; Flag Hill model with k <1/3 of Lowest Positive Dose and if this model is best, select next best model with p>0.05

1.2 Functional classifications parameters:

Gene Ontology Analyses: GO Categories:Biological process (or Select Pathway: REACTOME)
Remove BMD>highest dose from category descriptive statistics
Remove BMD with p-value < 0.1
Remove Genes with BMDU/BMDL>40
Eliminate Gene Set Redundancy
Identify Conflicting Probesets: correlation cutoff: 0.5
Most sensitive pathway was identified as pathway with lowest median BMD value when contained at least 3 genes that passed probe-level BMD filtering and when the proportion of these genes in the pathway reached at least 5%.

### 2. Supplemental tables

Supplemental Table 1 Comparison of methods used for hybridization of the raw Affymetrix Rat Genome 230 2.0 Array data in this study

| | Background estimation | Normalization | Summarization | Other |
|---|---|---|---|---|
| GCRMA | Model of PM and MM signals that include sequence-dependent non-specific binding and optical noise for each probe pair. | Quantile normalization | Robust multiarray linear model fit using Median Polish | -Employs MM intensities to correct background<br><br>-Results on log2 scale |
| RMA | Convolution model of the observed PM probe signal as a sum of signal and background. Multiplicative error. Single global background used to adjust all intensities. | Quantile normalization | Robust multiarray linear model fit using Median Polish | -Does not use MM intensities<br><br>-Results on log2 scale |
| MAS 5.0 | Weighted average of lowest 2% of signals from 16 equal rectangular regions. Weighting reflects distance of 16 averages from a particular probe.<br>PM signals adjusted by MM intensities to correct for non-specific binding.<br>Multiplicative error. | Linear scaling using the trimmed mean that excludes the highest and lowest 2% of the data | Robust single-array method using one step Tukey biweight of background corrected intensities | -Employs MM intensities<br><br>-Results on linear scale |
| MAS 5.0_noA | As in MAS 5.0 | As in MAS 5.0 | As in MAS 5.0 | -Probes with only “Absent” calls across all specimens are removed<br><br>-Results on linear scale |
| PLIER* | PM-MM method. Mixed error model. | Quantile normalization | Multiplicative model fitted to the PM-MM values, with array and probe as effects. An M-estimator is used that assess goodness of fit by the residuals generated from a generalized log transformation based on the PM and MM | -Employs PM and MM intensities.<br><br>-Results on linear scale |
| PLIER16* | As in PLIER | As in PLIER | As in PLIER | -As in PLIER<br>-16 is added to normalized signal intensities |
| PLIER16_noA* | As in PLIER16 | As in PLIER16 | As in PLIER16 | -As in PLIER16<br><br>-Probes with only “Absent” calls across all specimens are removed |

* Relates to PLIER as specifically implemented in this study

Supplemental table 2 Median lowest BMD (BEPOD) and BMDL (BEPOD/L) values [mg/kg-day] determined by toxicogenomic BMD modeling for different normalization methods and different gene sets

| Collection of gene sets | **GCRMA** | **RMA** | **MAS 5.0** | **MAS5.0_ _noA calls** | **PLIER** | **PLIER16** | **Plier16_ _no A calls** |
|---|---|---|---|---|---|---|---|
| **Crude MCMH** | | | | | | | |
| GO:BP | | | | | | | |
| BMD | 79.34 | 69.48 | 62.67 | 62.67 | 61.69 | 61.69 | 61.69 |
| BMDL | 49.80 | 43.77 | 41.86 | 41.86 | 41.36 | 41.36 | 41.36 |
| Reactome | | | | | | | |
| BMD | 79.36 | 81.32 | 71.88 | 71.86 | 79.21 | 70.21 | 79.21 |
| BMDL | 49.80 | 50.42 | 47.15 | 47.15 | 49.59 | 49.59 | 49.59 |
| **MCMH** | | | | | | | |
| GO:BP | | | | | | | |
| BMD | 69.45 | 79.21 | 58.84 | 58.84 | 95.95 | 95.95 | 95.95 |
| BMDL | 46.21 | 49.59 | 40.76 | 40.76 | 65.58 | 65.58 | 65.58 |
| Reactome | | | | | | | |
| BMD | 223.52 | 284.71 | 174.02 | 174.02 | 271.37 | 264.09 | 264.09 |
| BMDL | 155.45 | 186.10 | 118.79 | 118.79 | 172.32 | 172.40 | 172.40 |
| **p-toluidine** | | | | | | | |
| GO:BP | | | | | | | |
| BMD | 2.88 | 3.00 | 0.33 | 0.33 | 2.82 | 5.29 | 5.29 |
| BMDL | 1.12 | 1.12 | 0.07 | 0.07 | 1.01 | 2.13 | 2.13 |
| Reactome | | | | | | | |
| BMD | 5.32 | 8.09 | 3.33 | 5.83 | 2.85 | 15.77 | 15.77 |
| BMDL | 1.73 | 4.89 | 0.66 | 2.52 | 0.98 | 10.99 | 10.99 |
| **DMPT - liver** | | | | | | | |
| GO:BP | | | | | | | |
| BMD | 1.47 | 1.72 | 1.04 | 1.04 | 1.02 | 1.36 | 1.36 |
| BMDL | 0.50 | 0.96 | 0.34 | 0.34 | 0.30 | 0.42 | 0.42 |
| Reactome | | | | | | | |
| BMD | 3.44 | 4.24 | 2.62 | 2.62 | 2.43 | 5.17 | 6.62 |
| BMDL | 1.79 | 2.05 | 1.10 | 1.10 | 0.94 | 1.59 | 3.04 |
| **PPH-kidney** | | | | | | | |
| GO:BP | | | | | | | |
| BMD | 22.65 | 53.35 | 63.40 | 63.40 | 118.53 | 176.02 | 176.02 |
| BMDL | 4.20 | 16.40 | 11.26 | 11.26 | 75.35 | 120.34 | 120.34 |
| Reactome | | | | | | | |
| BMD | 54.51 | 183.97 | 243.13 | 243.13 | 201.67 | 180.81 | 180.81 |
| BMDL | 8.58 | 120.87 | 104.12 | 152.92 | 116.43 | 120.34 | 120.34 |

Supplemental table 3: GO:BP ontologies and Reactome Pathways with lowest median BMD or BMDL

| Accession code | Name |
|---|---|
| GO:0002933 | Lipid hydroxylation |
| GO:0006084 | Acetyl-CoA metabolic process |
| GO:0006090 | Pyruvate metabolic process |
| GO:0007040 | Lysosome organization |
| GO:0007062 | Sister chromatid cohesion |
| GO:0009108 | Coenzyme biosynthetic process |
| GO:0032026 | Response to magnesium ion |
| GO:0034502 | Protein localization to chromosome |
| GO:0035384 | Thioester biosynthetic process |
| GO:0042130 | Negative regulation of T cell proliferation |
| GO:0043588 | Skin development |
| GO:0043651 | Linoleic acid metabolic process |
| GO:0044246 | Regulation of multicellular organismal metabolic process |
| GO:0048266 | Behavioral response to pain |
| GO:0051445 | Regulation of meiotic cell cycle |
| GO:0061051 | Positive regulation of cell growth involved in cardiac muscle cell development |
| GO:0070141 | Response to UV-A |
| GO:0070206 | Protein trimerization |
| GO:0070207 | Protein homotrimerization |
| GO:0070988 | Demethylation |
| GO:0070989 | Oxidative demethylation |
| GO:0071616 | Acyl-CoA biosynthetic process |
| GO:0080171 | Lytic vacuole organization |
| GO:0090207 | Regulation of triglyceride metabolic process |
| GO:1902229 | Regulation of intrinsic apoptotic signaling pathway in response to DNA damage |
| GO:1904705 | Regulation of vascular smooth muscle cell proliferation |
| GO:2000727 | Positive regulation of cardiac muscle cell differentiation |
| R-RNO-1296072 | Voltage gated Potassium channels |
| R-RNO-141424 | Amplification of signal from the kinetochores |
| R-RNO-141444 | Amplification of signal from unattached kinetochores via a MAD2 inhibitory signal |
| R-RNO-1483206 | Glycerophospholipid biosynthesis |
| R-RNO-156580 | Phase II - Conjugation of compounds |
| R-RNO-159418 | Recycling of bile acids and salts |
| R-RNO-163125 | Post-translational modification: synthesis of GPI-anchored proteins |
| R-RNO-163685 | Integration of energy metabolism |
| R-RNO-1638091 | Heparan sulfate/heparin (HS-GAG) metabolism |
| R-RNO-174154 | APC/C:Cdc20 mediated degradation of Securin |
| R-RNO-174154 | APC/C:Cdc20 mediated degradation of Securin |
| R-RNO-174178 | APC/C:Cdh1 mediated degradation of Cdc20 and other APC/C:Cdh1 targeted proteins in late mitosis/early G1 |
| R-RNO-174184 | Cdc20:Phospho-APC/C mediated degradation of Cyclin A |
| R-RNO-176417 | Phosphorylation of Emi1 |
| R-RNO-179419 | APC:Cdc20 mediated degradation of cell cycle proteins prior to satisfation of the cell cycle checkpoint |
| R-RNO-2024096 | HS-GAG degradation |
| R-RNO-2187338 | Visual phototransduction |
| R-RNO-382556 | ABC-family proteins mediated transport |
| R-RNO-5173105 | O-linked glycosylation |
| R-RNO-5663220 | RHO GTPases Activate Formins |
| R-RNO-69618 | Mitotic Spindle Checkpoint |

| Accession code | Name |
|---|---|
| R-RNO-75105 | Fatty acyl-CoA biosynthesis |
| R-RNO-75105 | Fatty acyl-CoA biosynthesis |
| R-RNO-77289 | Mitochondrial Fatty Acid Beta-Oxidation |
| R-RNO-8878166 | Transcriptional regulation by RUNX2 |
| R-RNO-8978868 | Fatty acid metabolism |

### 3. Supplemental figures

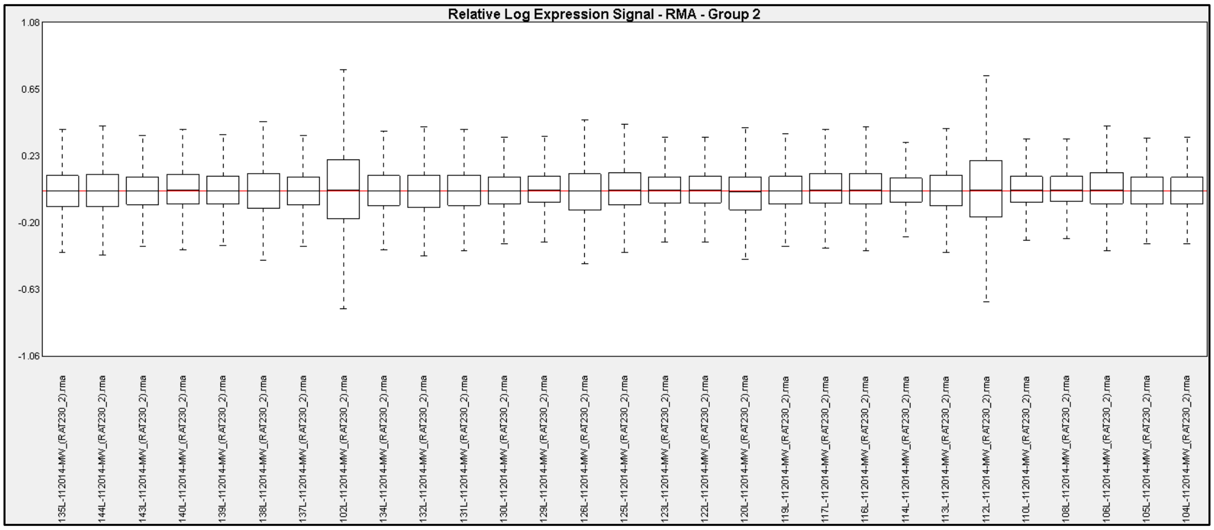


Supplemental figure 1  RLE boxplots of expression data for crude MCHM normalized by RMA before removing low-quality chips 102L and 112 L.

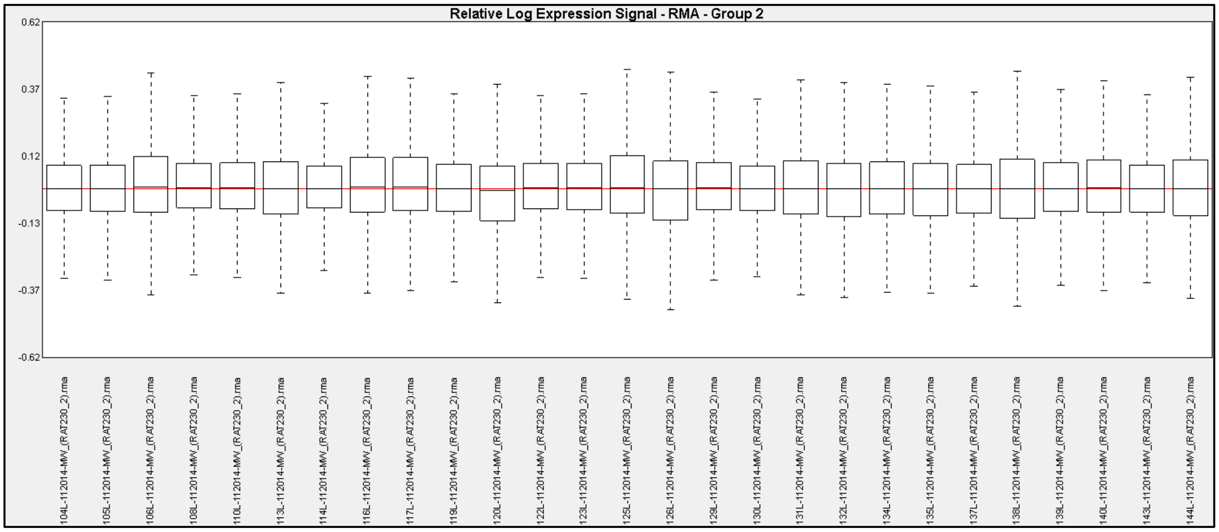


Supplemental figure 2  RLE boxplots of expression data for crude MCHM normalized by RMA after removing low-quality chips 102L and 112 L.

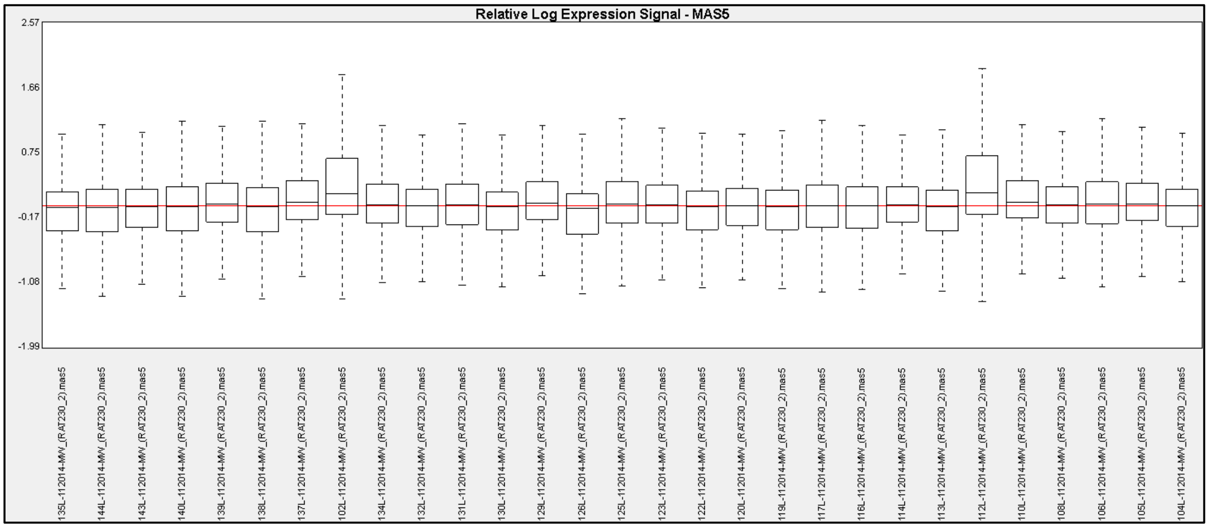


Supplemental figure 3 RLE boxplots of expression data for crude MCHM normalized by MAS 5.0 method before removing low-quality chips 102L and 112 L.

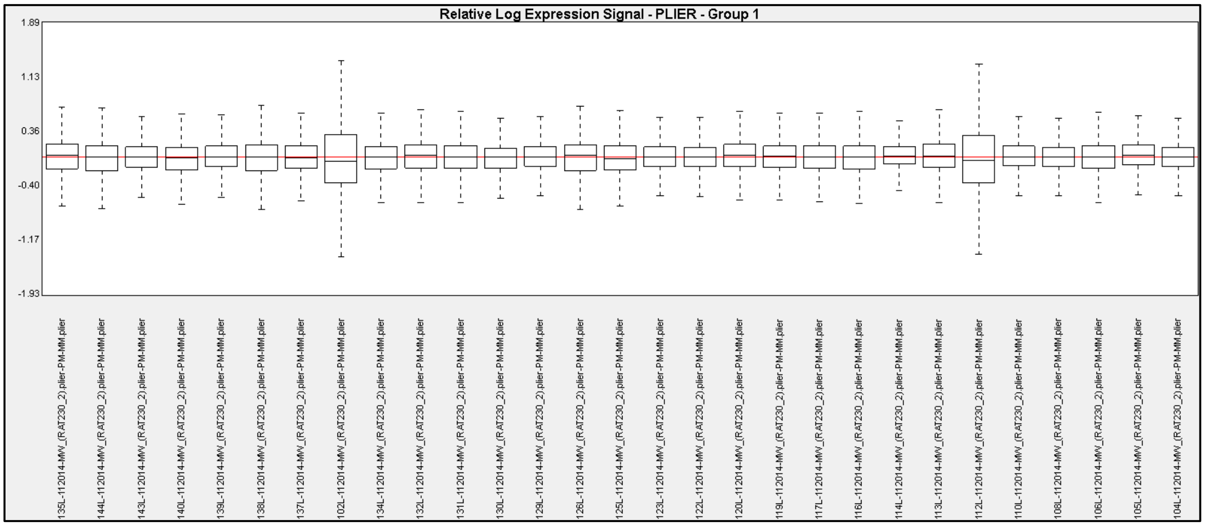


Supplemental figure 4 RLE boxplots of expression data for crude MCHM normalized by PLIER before removing low-quality chips 102L and 112 L.

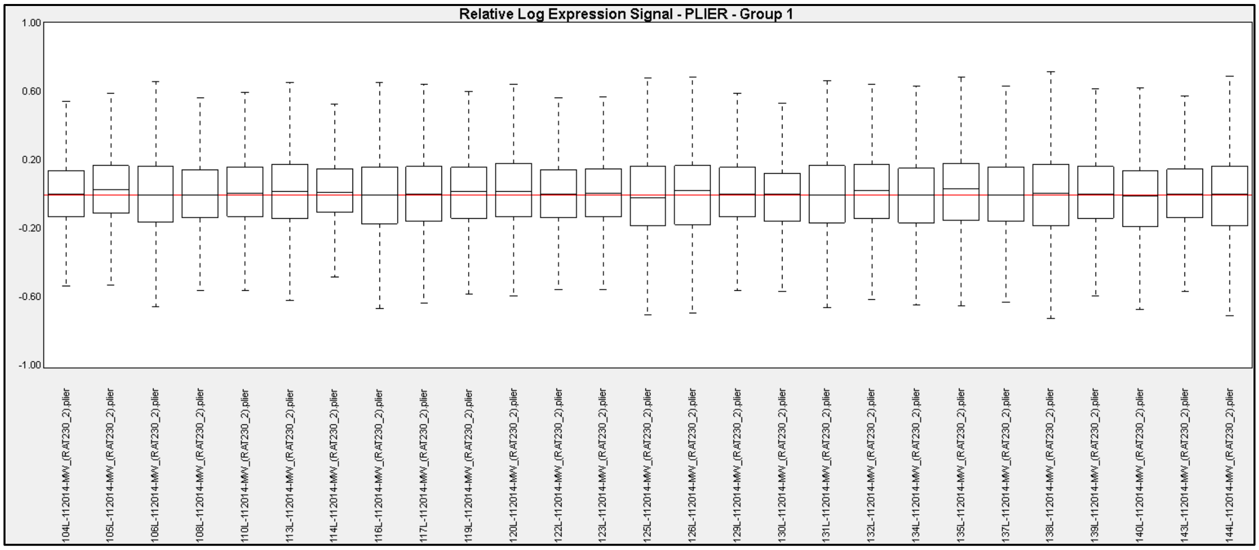


Supplemental figure 5    RLE boxplots of expression data for crude MCHM normalized by PLIER after removing low-quality chips 102L and 112 L.

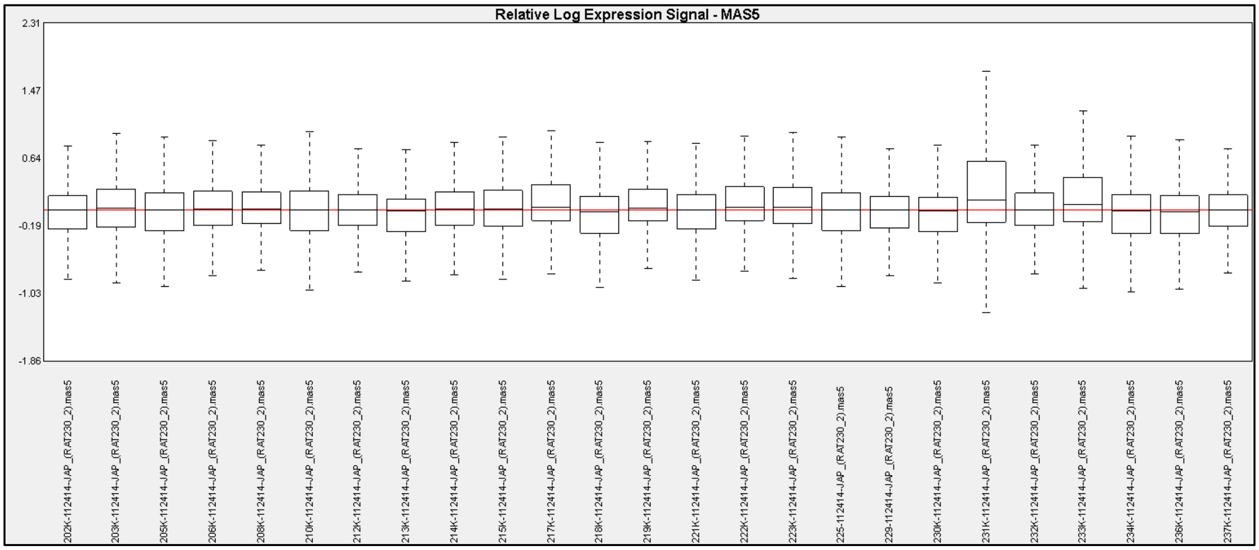


Supplemental figure 6    RLE boxplots of expression data for PPH normalized by MAS 5.0 before removing low-quality chip 231K.

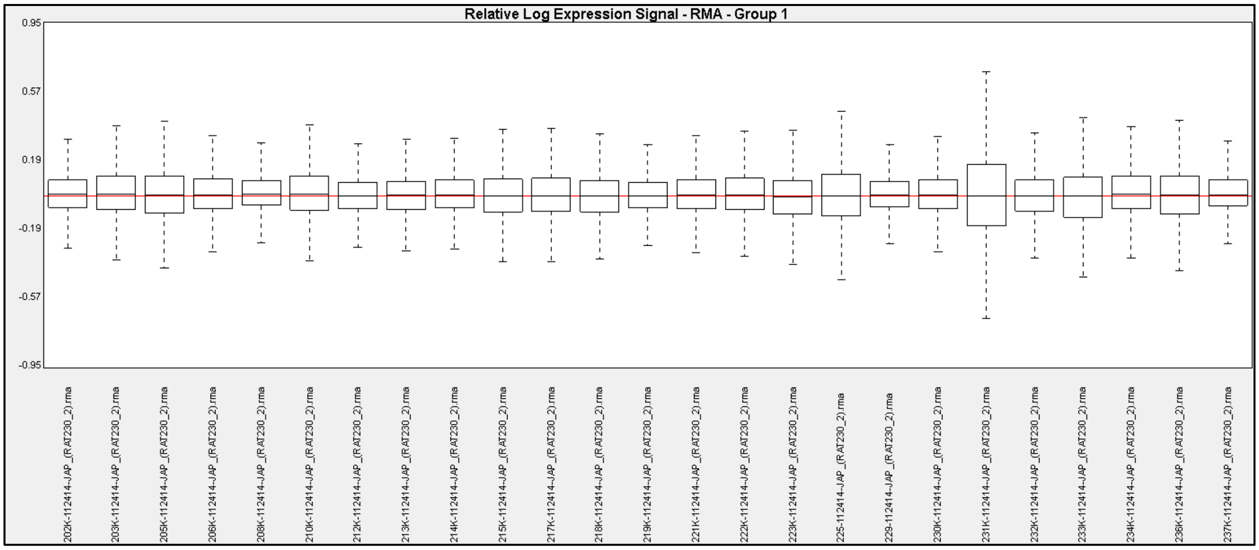


Supplemental figure 7 RLE boxplots of expression data for PPH normalized by RMA before removing low-quality chip 231K.

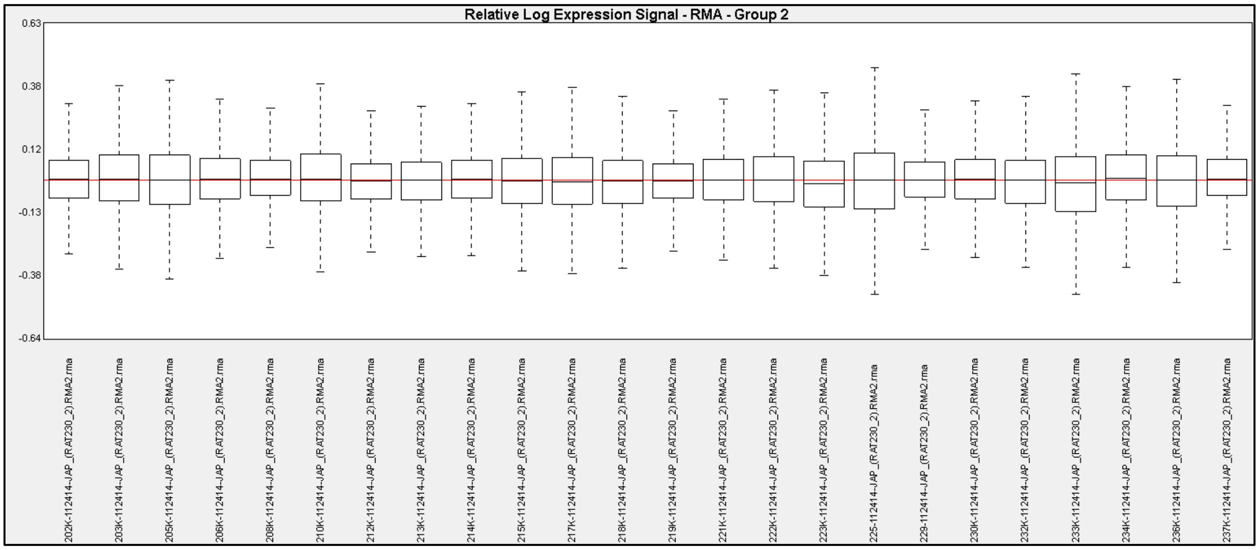


Supplemental figure 8 RLE boxplots of expression data for PPH normalized by RMA after removing low-quality chip 231K.

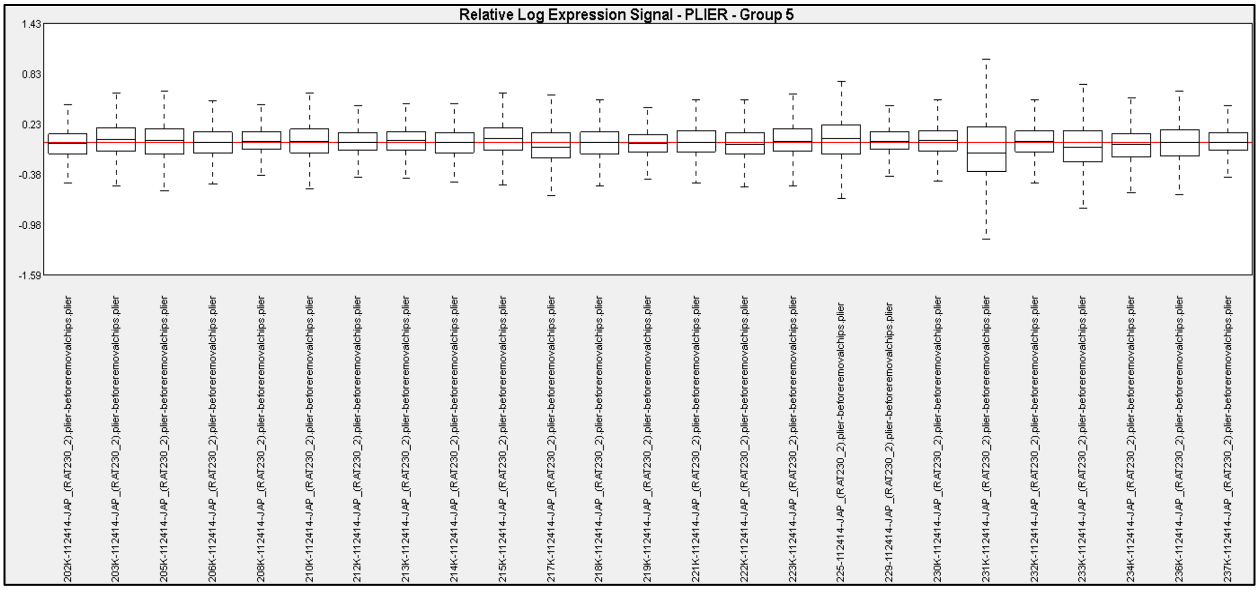


Supplemental figure 9 RLE boxplots of expression data for PPH normalized by PLIER before removing low-quality chips 217K, 225K, 231K and 233K.

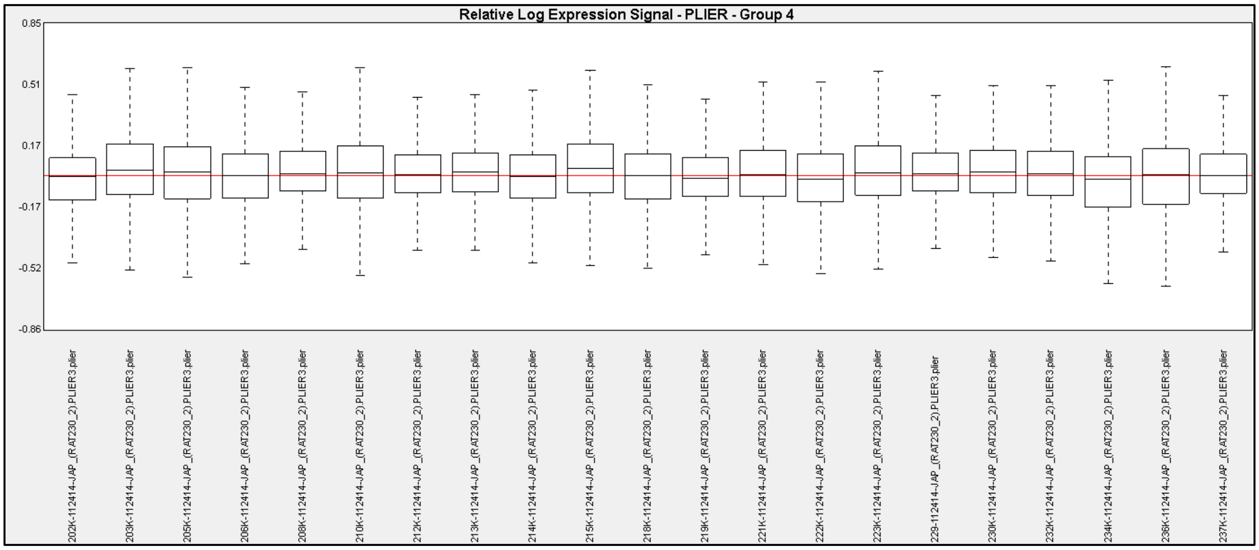


Supplemental figure 10 RLE boxplots of expression data for PPH normalized by PLIER after removing low-quality chips 217K, 225K, 231K and 233K.

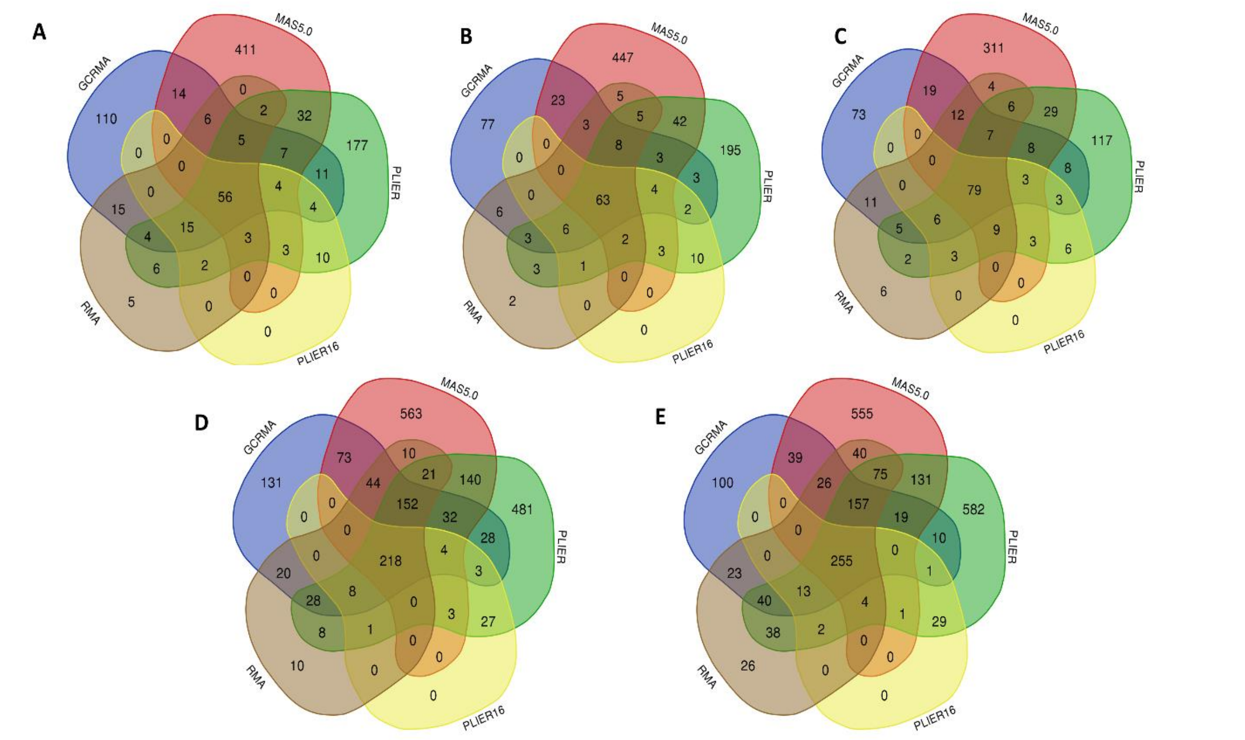


Supplemental figure 11 Numbers of differentially expressed genes identified for chemical-treatment pairs by different normalization methods depicted by Venn diagrams. A – crude MCHM/liver; B- neat MCHM/liver; C – PPH/kidney; D – DMPT/liver; E – p-toluidine/liver. Only GCRMA, RMA, MAS5.0, PLIER and PLIER16 are shown to enable visualization (normalization methods MAS5.0_noA and PLIER16_noA identify subsets of DEG found by MAS5.0 and PLIER16, respectively).

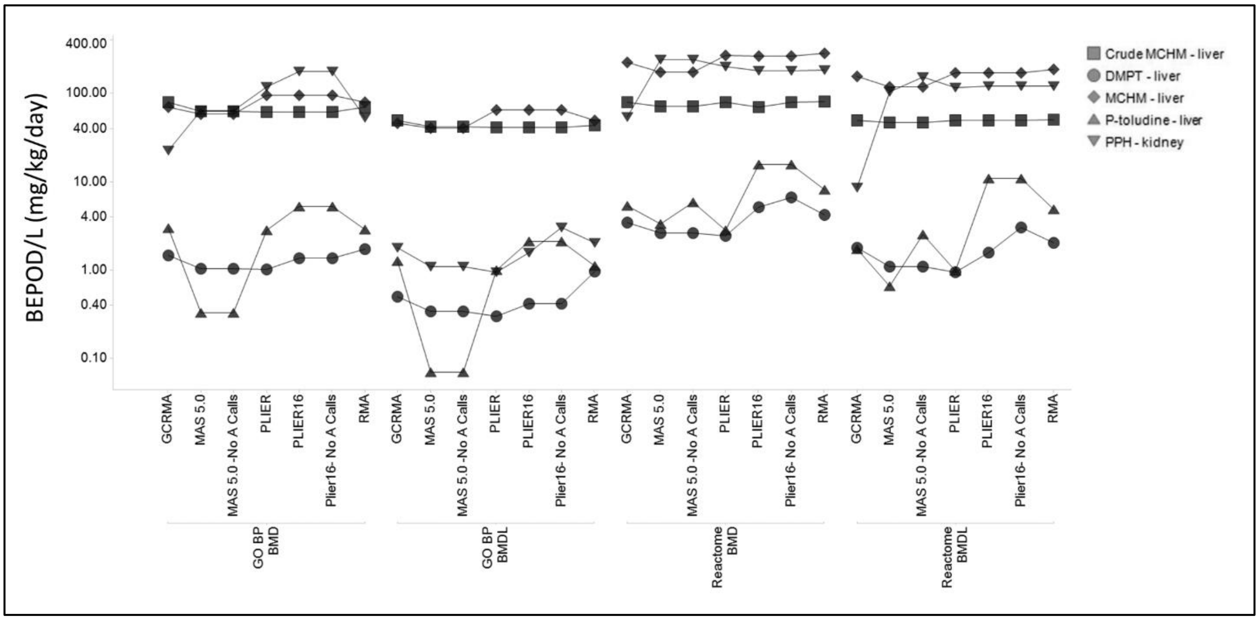


Supplemental figure 12 Lowest median BMD (BEPOD) and BMDL (BEPOD/L) values for different gene sets (GO:BP or Reactome) and normalizations (GCRMA, MAS5.0, MAS5.0_noA calls, PLIER, PLIER16, PLIER16_noA calls and RMA).
