## Supplementary material for "The sensitivity of transcriptomics BMD modeling to the methods used for microarray data normalization": BMDExpress 2.2 Project File: Mezencev_Auerbach_BMDExpress_ProjectFile_readme.pdf

### BMDExpress 2.0 Project File

Dataset labeling

Each expression dataset is labelled according to the following scheme:

**Chemical_tissue type_normalization method_bmd**

Example: “crude_MCHM_liver_GCRMA_bmd” dataset represents expression data from GeneChip Rat Genome 230 2.0 Array for livers of rats exposed to crude MCHM and normalized by GCRMA method.

These data were generated from public data available in Gene Expression Omnibus (GEO) repository as series with accession numbers GSE75655, GSE75657, GSE75656, and GSE100502.

Abbreviations of chemical names and pre-processing (“normalization”) methods:

crude 4-methylcyclohexanemethanol (crude-MCHM)
neat 4-methylcyclohexanemethanol (MCHM)
propylene glycol phenyl ether (PPH)
N,N-dimethyl-p-toluidine (DMPT)
p-toluidine

MAS5.0.0 (Microarray Affymetrix Suite version 5.0)
RMA (Robust Multichip Analysis)
GCRMA (GeneChip Robust Multichip Analysis)
PLIER (Probe Logarithmic Intensity Error Estimation)
PLIER16: value of 16 was added to PLIER-normalized expression values
MAS5.0_noA and PLIER16_noA: Datasets with removed probesets that showed "Absent" MAS5.0 absolute detection calls across all specimens from corresponding MAS5.0 and PLIER16-normalized data

Columns in datasets

Description of data structure in BMDExpress 2.2 files is available at https://github.com/auerbachs/BMDExpress-2/wiki
